# DNA-uptake pilus of *Vibrio cholerae* capable of kin-discriminated auto-aggregation

**DOI:** 10.1101/354878

**Authors:** David. W. Adams, Sandrine Stutzmann, Candice Stoudmann, Melanie Blokesch

**Affiliations:** Laboratory of Molecular Microbiology, Global Health Institute, School of Life Sciences, Station 19, EPFL-SV-UPBLO, Swiss Federal Institute of Technology Lausanne (Ecole Polytechnique Fédérale de Lausanne; EPFL), CH-1015 Lausanne, Switzerland.

**Keywords:** Auto-aggregation, DNA uptake, natural transformation, pilus dynamics, type IV pilus, *Vibrio cholerae*

## Abstract

Natural competence for transformation is a widely used and key mode of horizontal gene transfer that can foster rapid bacterial evolution. Competent bacteria take-up DNA from their environment using Type IV pili, a widespread and multi-purpose class of cell surface polymers. However, how pili facilitate DNA-uptake has remained unclear. Here, using direct labelling, we show that in the Gram-negative pathogen *Vibrio cholerae* DNA-uptake pili are highly dynamic and that they retract prior to DNA-uptake. Unexpectedly, these pili can self-interact to mediate auto-aggregation of cells into macroscopic structures. This phenotype is conserved in disease causing pandemic strains. However, extensive strain-to-strain variability in the major pilin subunit PilA, present in environmental isolates, controls the ability of pili to interact without affecting transformation. We go on to show that interactions between pili are highly specific, enabling cells producing pili composed of different PilA subunits to discriminate between one another. On chitin surfaces, a natural habitat of *V. cholerae*, pili connect cells within dense networks, suggesting a model whereby DNA-uptake pili function to promote inter-bacterial interactions during surface colonisation. Moreover, our results provide evidence that type IV pili could provide a simple and potentially widespread mechanism for bacterial kin recognition.

## Introduction

How bacteria physically sense and interact with their environment is a fundamental problem in biology. Type IV pili (T4P) are cell surface polymers ideally suited to this task^1,2^. Composed of a single major pilin and assembled by widely distributed and conserved machinery, T4P exhibit extensive functional versatility, with roles in motility, DNA-uptake, surface sensing and adhesion^3,5^. Consequently, T4P are critical virulence factors for a number of important human pathogens including *Vibrio cholerae*, which causes the pandemic diarrhoeal disease cholera^6^. In Gram-negative bacteria pilins are processed at the inner-membrane, extracted by the assembly machinery and polymerised into a helical pilus fibre that exits the cell surface through a gated outer-membrane pore; the secretin^7,11^. A key feature of T4P is their ability to undergo dynamic cycles of extension and retraction^12,13^, powered by the action of dedicated extension (*e.g.* PilB) and retraction (*e.g.* PilT) ATPases, which either add or liberate pilin subunits at the base^14,15^. These dynamics are essential for many T4P functions *e.g.* twitching-motility^11,12^. Thus, understanding how T4P function may yield insights valuable for understanding mechanisms of environmental survival and pathogenesis.

Despite their multifunctional potential, pandemic strains of *V. cholerae* typically encode three distinct T4P systems – two of which are well characterised. First, toxin co- regulated pili (TCP) serve a dual role as both a receptor for CTXϕ bacteriophage^16^, which carries the cholera toxin genes, and as the primary human colonisation factor with multiple essential roles in infection involving adhesion and auto-aggregation on the intestinal cell surface^17,19^. Second, Mannose-sensitive haemagglutinin (MSHA) pili are involved in surface sensing and attachment and thus, are important in the initiation of biofilm formation^20,24^. Third, in its natural aquatic environment *V. cholerae* often associates with chitinous surfaces^25^, which are nutritious, foster biofilm formation and likely play a role in environmental dissemination and transmission to humans^26^-^29^. Chitin utilisation triggers competence for natural transformation^30^, a widely used mode of horizontal gene transfer that allows bacteria to take up DNA from their environment, and which can thus, foster rapid bacterial evolution^31^. This requires the production of the Chitin-Chitin (ChiRP) or DNA-or DNA pilus^30^.

We previously showed DNA-previously showed DNA pili form *bona fide* pili composed of the major subunit PilA and that transformation was dependent on the presumed retraction ATPase PilT^32^. However, the pilus itself is not sufficient for transformation and requires the concerted action of a periplasmic DNA-binding protein, ComEA^32,33^. Upon receipt of transforming DNA ComEA switches from a diffuse to focal localisation^33,34^. These findings, together with work in other organisms, led to a model in which pilus retraction facilitates DNA entry into the periplasm^35^, wherein ComEA acts as ‘ratchet’ to pull in the remaining DNA^33^. Subsequently, DNA transport across the inner-membrane occurs via a spatially coupled channel, ComEC^34,36^. Though this model is well supported by genetic experiments^32^ and the similarly combined action of T4P and ComEA in other organisms^37,38^, direct evidence is lacking.

Here, we visualised the DNA-uptake pilus directly using a cysteine labelling approach, which was recently validated as a tool for labelling pili^39^. As predicted, we demonstrate that the pili are highly dynamic and that these dynamics are PilT-dependent. Unexpectedly, however, we discovered that DNA-uptake pili are also capable of self-interacting, which in liquid culture results in a strong auto-aggregation phenotype. Variability in PilA controls this activity but has no affect on transformation. Remarkably, specific interactions allow pili composed of different PilA subunits to distinguish between one another, enabling a simple mechanism for kin recognition.

## Results

### Direct observation of pilus dynamics by cysteine labelling

To avoid the limitations imposed by immuno-fluorescent methods, we employed a cysteine labelling approach using a thiol-reactive dye^39,40^. PilA cysteine variants were created along the length of the surface exposed aβ-loop^9,11^ and tested for functionality using a previously validated chitin-independent transformation system in which competence induction is arabinose-inducible^41^ (Tn*tfoX*; see methods) (Fig. 1A, B and Fig. S1A). A variant, PilA[S67C], was identified that relative to the unmodified parent is fully transformable (Fig.1A), produced at similar levels (Fig. S1B), and does not affect pilus assembly (Fig. S1C), as assayed by a classical shearing method. Since other variants exhibited functional defects they were not studied further.

**Figure 1.**
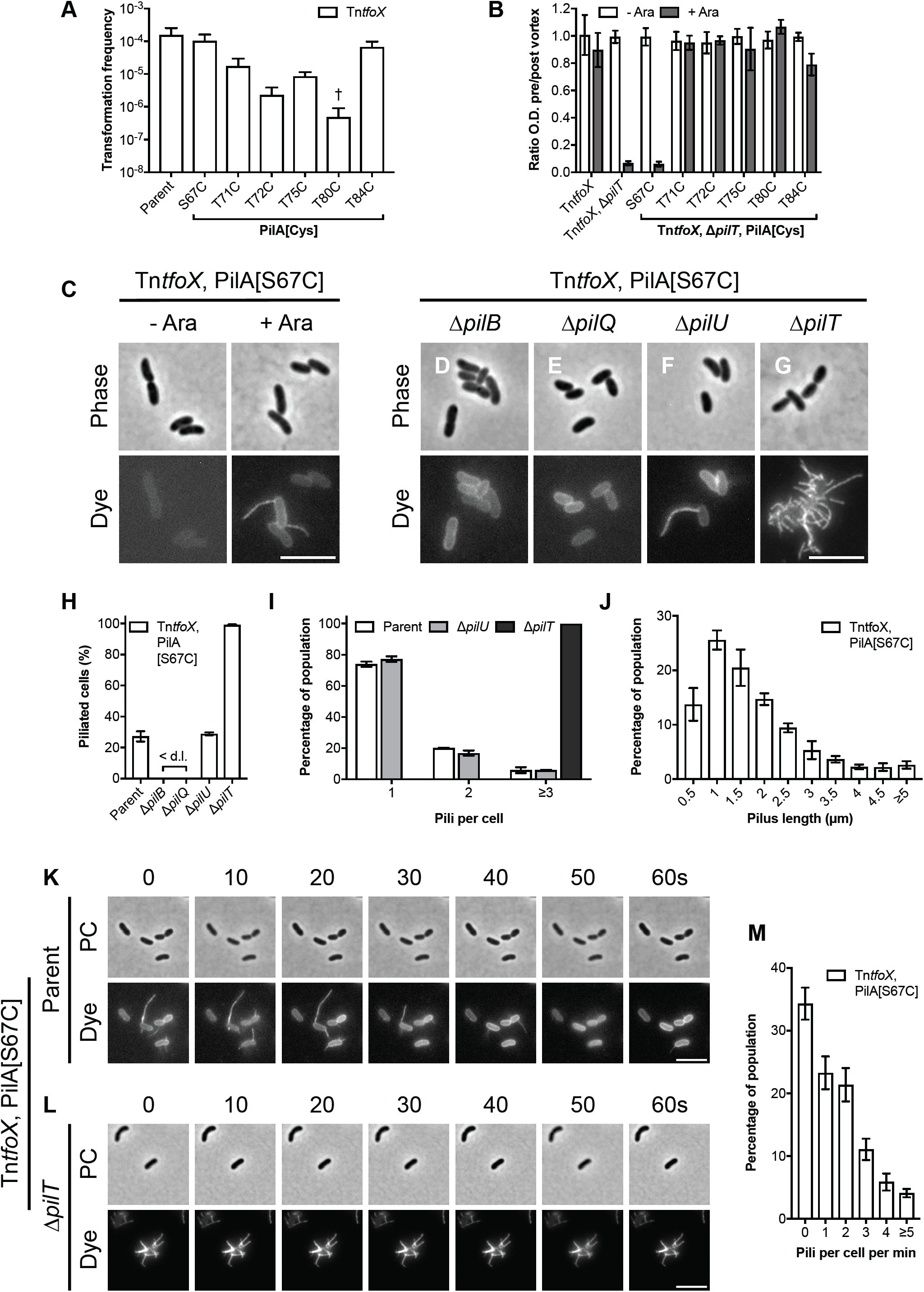
Direct observation of dynamic DNA-uptake pili. **(A)** Functionality of PilA cysteine variants (PilA[Cys]) in a chitin-independent transformation assay using strains carrying an arabinose-inducible copy of *tfoX* (Tn*tfoX*). Transformation frequencies are the mean of three independent biological repeats (+S.D.). < d.l., below detection limit. †, < d.l. in one repeat. **(B)** Effect of PilA cysteine variants on the ability of retraction deficient (Tn*tfoX*, Δ*pilT*) cells to aggregate. Aggregation is shown as the ratio of the culture optical density (O.D. _600_) before and after vortexing, in the absence (- Ara) and presence (+ Ara) of *tfoX* induction, as indicated. Values are the mean of three independent biological repeats (±S.D.). **(C-G)** Snapshot imaging of pili in cells of A1552-Tn*tfoX*, PilA[S67C] and its derivatives. **(C)** Cells of A1552-Tn*tfoX*, PilA[S67C] were grown in the absence (- Ara) and presence (+ Ara) of *tfoX* induction, as indicated, and stained with AF-488-Mal. Bar = 5 μm. (D-E) Cells of A1552-Tn*tfoX*, PilA[S67C] were grown in the presence of *tfoX* induction and stained with AF-488-Mal in a **(D)** Δ*pilB*, **(E)** Δ*pilQ*, **(F)** Δ*pilU* and **(G)** Δ*pilT* background, as indicated. Bar = 5 μm. **(H-J)** Quantification of piliation in snapshot imaging. Bars represent the mean of three repeats (±S.D.). **(H)** Percentage of piliated cells in the indicated backgrounds. *n* = *c.a.* 2000 cells per strain per repeat. **(I)** Histogram of number of pili per cell in piliated cells in WT parent (A1552-Tn*tfoX*, PilA[S67C]), Δ*pilU* (A1552-Tn*tfoX*, PilA[S67C], Δ*pilU*) and Δ*pilT* (A1552-Tn*tfoX*, PilA[S67C], Δ*pilT*) backgrounds, as indicated. *n* = *c.a.* 300 cells per strain per repeat. **(J)** Histogram of pilus length in cells of A1552-Tn*tfoX*, PilA[S67C]. *n* = *c.a.* 500-600 cells per repeat. **(K-L)** Time-lapse series of pilus dynamics in **(K)** WT parent (A1552-Tn*tfoX*, PilA[S67C]) and **(L)** Δ*pilT* (A1552-Tn*tfoX*, PilA[S67C], Δ*pilT*) backgrounds. Cells were stained with AF-488-Mal and imaged at 10s intervals for 1 minute. Upper panels show phase-contrast (PC), lower panels show fluorescence (dye). Time in seconds (s), as indicated. Bars = 5 μm. **(M)** Histogram showing quantification of pili produced per cell per min in the WT parent (A1552-Tn*tfoX*, PilA[S67C]) background. Bars represent the mean of three repeats (±S.D.). *n* = *c.a.* 500-700 cells per repeat.

When stained, competent cells producing PilA[S67C] exhibited visible pili (Fig. 1C). On average 27 ± 3 % cells were piliated, with the majority of these displaying 1 or 2 pili per cell (Fig. 1H and I). The length of pili was clustered around 1-2 μm, though pili up to 10 μm in length were also observed (Fig. 1J). Numerous detached pili were also evident in the growth media. Intriguingly, these pili frequently appeared to self-interact, forming large structures composed of networks of pili (Fig. S2A). When examined by time-lapse microscopy cells exhibited rapid pilus dynamics, with multiple assembly and retraction events immediately apparent (Fig. 1K; Movie S1-4). Indeed, within the 1 minute time frame studied 66 ± 3 % cells exhibited pili, with most cells producing 1-2 pili per minute (Fig. 1M). Notably, a small subpopulation of cells was even more dynamic, elaborating ≥5 pili per minute. Consistent with this dynamic behaviour, and in support of the hypothesis that pilus retraction precedes DNA-uptake, when cells were provided with purified genomic DNA it was possible to concurrently visualise pilus retraction followed by DNA-uptake, as monitored by the re-localisation of ComEA-mCherry, which binds incoming DNA in the periplasm (Fig. S3; Movie S5).

As expected, deletion of components required for pilus assembly (*e.g.* the assembly ATPase PilB or the secretin PilQ) abolished piliation (Fig. 1D and E). However, despite the absence of obvious pili the cell body still stained. Control experiments in which PilA[S67C], but not PilA[WT], was produced independently of the normal assembly machinery also stained similarly (Fig. S1D), suggesting that this effect results from the dye being taken up non-specifically and retained by the inner-membrane pool of PilA[S67C].

As in other species *V. cholerae* encodes two potential retraction ATPases, PilT and PilU. Deletion of *pilU* did not affect piliation (Fig. 1F, H and I), consistent with its dispensability for transformation^32^. In contrast, cells lacking the presumed retraction ATPase PilT were hyper-piliated, with essentially all cells displaying multiple static pili (Fig. G-I; Movie S6-7). Taken together with the dynamics described above, these data are consistent with the presence of multiple assembly complexes scattered across the cell, as previously predicted based on the mobility of PilB and the presence of multiple PilQ foci^32,42^. This might normally serve to facilitate rapid switching of pilus location or else might reflect a need to produce multiple pili under certain conditions. Unexpectedly, cells blocked for retraction were often grouped into small clusters within dense networks of pili as well as occasionally large aggregates of pili-encased cells (Fig. S2B and C). Indeed, when grown in liquid culture cells appeared to auto-aggregate.

### Competent cells auto-aggregate in the absence of pilus retraction

To test directly whether the aggregation phenotype was an inherent property of hyper-piliated cells or merely an artefact introduced by the cysteine variant, cells containing unmodified *pilA* were examined (Fig. 2A-C). Remarkably, aggregation occurred specifically in competence-induced cells lacking *pilT*, with the formation of large, multi-layered spherical aggregates on the order of 25-100 μm, which went on to form macroscopic aggregates that rapidly sediment (Fig. 2B, +Ara). Quantification of the ratio of cells remaining in solution to those in the settled aggregates (Fig. 2D) revealed that upon competence induction, ≥90% of the retraction deficient cells were present in aggregates (Fig. 2D). Notably, strains producing the PilA[S67C] variant used for labelling behave similarly (Fig. 1B).

**Figure 2.**
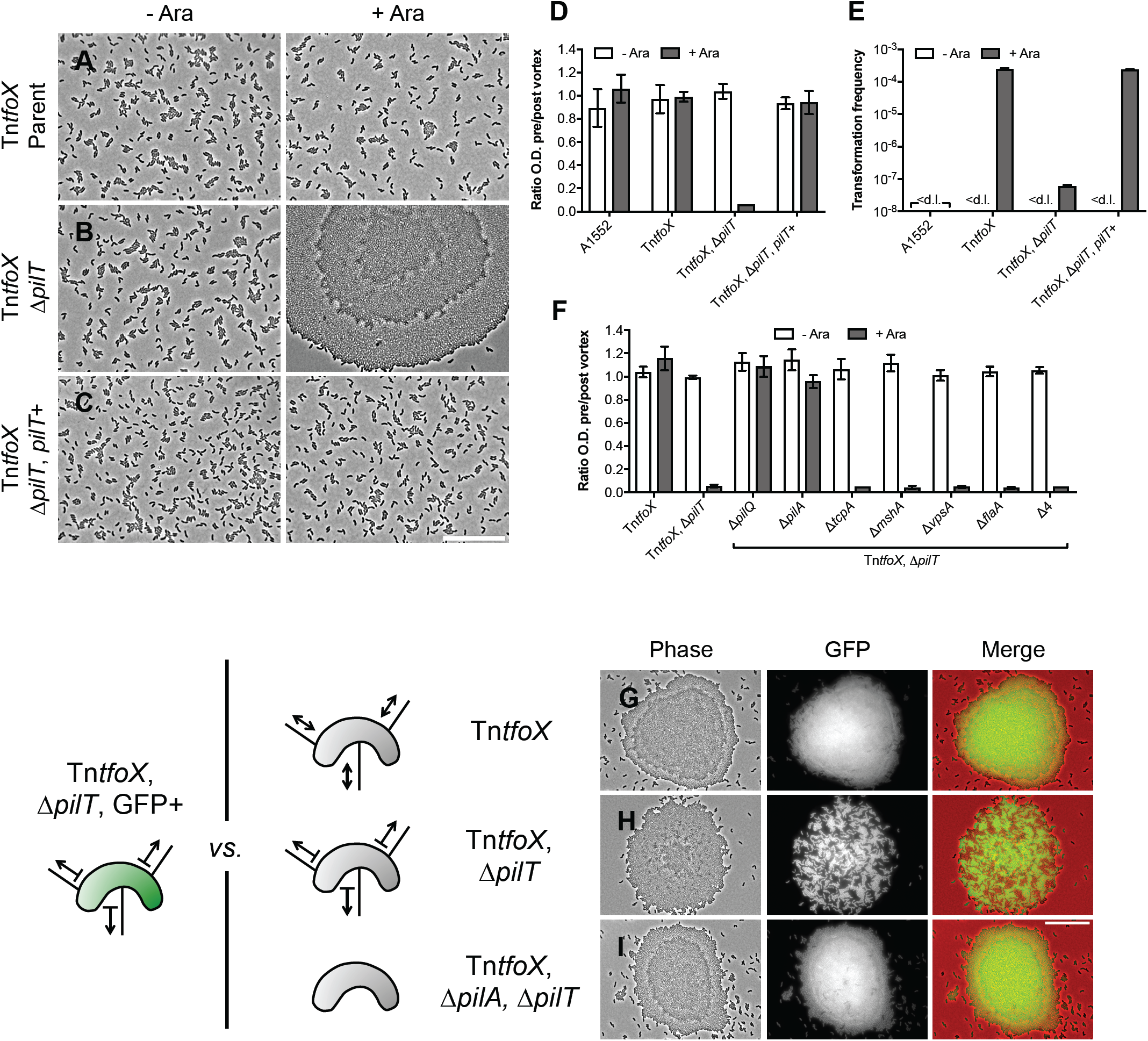
Competent cells auto-aggregate in the absence of pilus retraction. **(A-C)** Phase-contrast microscopy of cells of strains **(A)** A1552-Tn*tfoX*, **(B)** A1552-Tn*tfoX*, Δ*pilT* and (**C**) A1552-Tn*tfoX*, Δ*pilT, pilT*+, grown in the absence (- Ara) and presence (+ Ara) of *tfoX* induction, as indicated. Scale bar = 25 μm. **(D-E)** Aggregation and transformation frequency of cells of strains A1552-Tn*tfoX*, A1552-Tn*tfoX*, Δ*pilT* and A1552-Tn*tfoX*, Δ*pilT, pilT*+, grown in the absence (- Ara) and presence (+ Ara) of *tfoX* induction, as indicated. A1552 (without inducible *tfoX*) was used as a negative control. **(D)** Aggregation is shown as the ratio of the culture optical density (O.D. _600_) before and after vortexing. Values are the mean of three repeats (±S.D.). **(E)** Chitin-independent transformation frequency assay. Values are the mean of three repeats (+S.D.). < d.l., below detection limit. **(F)** Effect of various deletion backgrounds on the ability of retraction deficient cells to aggregate. Aggregation is shown as the ratio of the culture optical density (O.D. _600_) before and after vortexing, in the absence (- Ara) and presence (+ Ara) of *tfoX* induction, as indicated. Δ4 = Δ*tcpA*, Δ*mshA*, Δ*vpsA*, Δ*flaA* quadruple mutant. Values are the mean of three repeats (±S.D.). **(G-I)** Co-culture of fluorescent Δ*pilT* cells (A1552-Tn*tfoX*, Δ*pilT*, GFP+) producing GFP and non-fluorescent cells of the **(G)** WT parent (A1552-Tn*tfoX*), **(H)** Δ*pilT* (A1552-Tn*tfoX*, Δ*pilT*) and **(I)** Δ*pilA*, Δ*pilT* (A1552-Tn*tfoX*, Δ*pilA*, Δ*pilT*), grown in the presence of *tfoX* induction. Merged images show GFP in green and phase-contrast in red. Bar = 25 μm.

To rule out the possibility that the deletion of *pilT* has either a polar or otherwise non-specific effect, an intact copy of *pilT* driven by its native promoter was placed at an ectopic locus. Importantly, complementation of Δ*pilT* was sufficient to fully restore the *~*1000-fold defect in transformation frequency and abolished the aggregation phenotype (Fig. 2D and E). Complementation also fully counteracted the enhanced motility phenotype of Δ*pilT* (Fig. S4), which occurs due to loss of function in the adhesive MSHA pilus and is in agreement with the previously established shared role of PilT in MSHA pilus function^24,43^.

Time-course experiments indicated that aggregation occurs abruptly, suggesting that rather than gradually increasing in size, large aggregates form via the accumulation of smaller ones (Fig. S5A and B). In support of this hypothesis, when two otherwise isogenic Tn*tfoX* Δ*pilT* strains, one of which constitutively produces GFP, were allowed to aggregate together they formed well-mixed aggregates (Fig. S5E). If, however, they were allowed to aggregate separately before being mixed (Fig. S5C and D), they formed large composite aggregates consisting of a patchwork of labelled and unlabelled cells (Fig. S5F). Finally, the aggregates remained stable and did not disperse, even after prolonged culture overnight.

### Aggregates form via pilus-pilus interactions

The data so far suggest that hyper-piliated cells auto-aggregate via their DNA-uptake pili. Indeed, deletions affecting DNA-uptake pilus assembly (*pilQ* and *pilA*) were sufficient to abolish aggregation (Fig. 2F). In contrast, deletions targeting the assembly of TCP (*tcpA*) and MSHA (*mshA*) pili, the *Vibrio* polysaccharide matrix (*vpsA*) required for biofilm formation^44^ or the flagellum (*flaA*), were dispensable for aggregation, both individually and in combination (Fig. 2F). Similarly, additional genes tested with roles in biofilm formation (*bap1, rbmA, rbmEF, vpsT*)^45,46^, adhesion (*gbpA*)^47^, cell shape (*crvA*)^48^ and an O-linked glycosylase (*VC0393* or *pglL*_*Vc*_)^49^ that could be involved in pilin modification all aggregated at levels indistinguishable from the unmodified parent (Fig. S6A). Finally, deletion of *VC0502*, an ‘orphan’ type IV pilin encoded on the *Vibrio* seventh pandemic island II, had no effect on transformation, aggregation or motility (Fig. S6B-D). Additional control experiments indicated that aggregation occurred similarly during growth under high salt conditions (*i.e.* LB-S 20 g/L NaCl), which are often used to better reflect the natural aquatic environment of *Vibrio sp*. (Fig. S6E). Likewise, the presence of Bovine serum albumin, which was previously reported to disrupt pilus-pilus interactions in *Neisseria gonorrhoeae*^50^, also had no effect (Fig. S6E). Curiously, however, we noticed that the strength of the aggregation phenotype is sensitive to the concentration of divalent cations (*e.g.* CaCl_2_ or MgCl_2_) (Fig. S6F), though the significance of this result remains unclear.

To investigate how the pilus mediates auto-aggregation *i.e* pilus-pilus, pilus-cell or otherwise, we co-cultured cells producing either dynamic pili (Tn*tfoX*), static pili (Tn*tfoX*, Δ*pilT*), or blocked for both retraction and pilus production (Tn*tfoX*, Δ*pilT*, Δ*pilA*) with a Tn*tfoX* Δ*pilT* strain constitutively producing GFP (Fig. 2G-I) and asked whether these cells were recruited into the GFP-labelled aggregates. Strikingly, intermixed aggregates were only observed when both cells were themselves capable of producing non-retractile pili, and hence aggregating (Fig. 2H), indicating that aggregates form via direct pilus-pilus interactions between DNA-uptake pili.

### MSHA pili do not mediate aggregation

MSHA pili are important for surface sensing and attachment during the switch to the sessile lifestyle that precedes biofilm formation, and pilus retraction is thought to be required for their function^20,24^. Since MSHA pili are constitutively produced, but retraction deficient cells only aggregate upon competence induction, the results above suggest that MSHA pili are unable to mediate auto-aggregation. To test this hypothesis directly we grew cells engineered to increase the levels of the second messenger c-di-GMP, which has been reported to enhance the assembly of MSHA pili on the cell surface^43^. Control experiments indicated that decreasing c-di-GMP levels cells did not induce aggregation (Fig. S7A and B). In contrast, under elevated c-di-GMP levels, cells formed large clumps that appeared similar to the aggregates described above and had a modest sedimentation phenotype (Fig. S7A and B). However, the assembly of these structures occurred independently of *pilT, pilA* and *mshA*, but was abolished by the loss of *vpsA* (Fig. S7C). Thus, we conclude that these structures are not aggregates held together by pili but rather biofilms held together by matrix polysaccharide.

### PilA from pandemic strains is sufficient for aggregation in a non-pandemic strain

A1552, the strain used throughout this work, is an O1 El Tor clinical isolate representative of the on-going 7^th^ cholera pandemic. To test whether the ability to aggregate is conserved among other pandemic strains we tested 6 additional O1 El Tor strains. In the absence of *pilT* all aggregated to similar levels as A1552, as might be expected given that pandemic strains are closely related (Fig. 3A). Surprisingly, however, N16961 did not aggregate unless its well-characterised *hapR* frame-shift mutation^51^, which renders it quorum-sensing (QS) defective, was first repaired (Fig. 3A). Since transcription of the genes required for pilus assembly are not regulated by QS^41^ this finding suggests that there may be an additional layer of post-transcriptional control.

**Figure 3.**
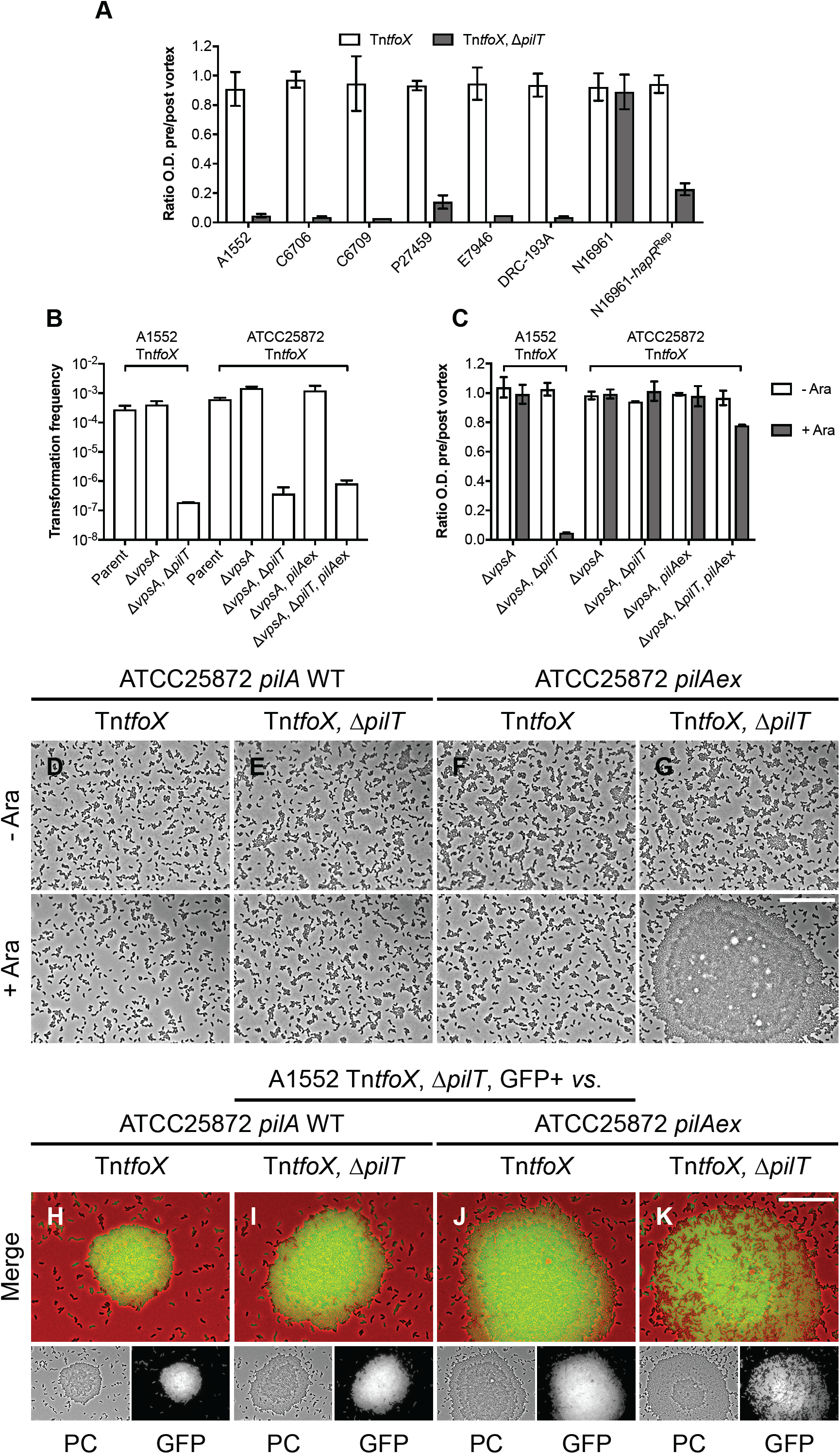
A1552 PilA is sufficient for aggregation in a non-pandemic strain. **(A)** Aggregation of representative 7^th^ pandemic strains of *V. cholerae*, including the effect of *hapR*^Rep^ on N16961, in a Tn*tfoX* and a Tn*tfoX*, Δ*pilT* background, as indicated. Aggregation is shown as the ratio of the culture optical density (O.D. _600_) before and after vortexing, in the presence of *tfoX* induction. Values are the mean of three repeats (±S.D.). **(B-C)** Transformation frequency and aggregation of *V. cholerae* strain ATCC25872-Tn*tfoX* compared to that of A1552-Tn*tfoX*. **(B)** Chitin-independent transformation assay. Transformation frequencies are the mean of three repeats (+S.D.). **(C)** Aggregation is shown as the ratio of the culture optical density (O.D. _600_) before and after vortexing, in the absence (- Ara) and presence (+ Ara) of *tfoX* induction, as indicated. Values are the mean of three repeats (±S.D.). **(D-G)** Phase-contrast microscopy of ATCC25872-Tn*tfoX*, Δ*vpsA* cells carrying (**D** and **E**) their native *pilA* (*pilA* WT) and (**F** and **G**) A1552 *pilA* (*pilA*ex), in a (**D** and **F**) *pilT*+ and (**E** and **G**) Δ*pilT* background, as indicated. Strains were cultured in the absence (- Ara) and presence (+ Ara), as indicated. Bar = 25 μm. Note that ATCC25872 derivatives were co-deleted for *vpsA* to rule out any compounding effects of biofilm formation. (**H**-**K**) Co-culture of fluorescent cells of A1552-Tn*tfoX*, Δ*pilT*, GFP+, producing GFP, and non-fluorescent cells of ATCC25872-Tn*tfoX*, Δ*vpsA* carrying (**H** and **I**) their native *pilA* (*pilA* WT) and (**J** and **K**) A1552 PilA (*pilA*ex), in a (**H** and **J**) *pilT*+ and (**I** and **K**) Δ*pilT* background, as indicated. Cells were grown in the presence of *tfoX* induction. Merged images show GFP in green and phase-contrast (PC) in red. Bar = 25 μm.

PilA is known to vary considerably between different environmental strains of *V. cholerae*^52^, whereas the majority of clinical isolates carry an identical *pilA*. Interesting exceptions are the toxigenic O37-serogroup strains V52 and ATCC25872, which were responsible for limited epidemics of a cholera-like illness in the 1960s^53,54^. These strains are thought to derive from a pandemic O1 classical strain through the exchange of the O-antigen cluster by horizontal gene transfer^55,56^. However, they both encode the same PilA version that is only 50% identical to that typically carried by pandemic strains (Fig. S8). Since V52 is QS deficient, and transformation is QS dependent, we investigated the functionality of this PilA in ATCC25872, which has a functional QS pathway. Indeed, ATCC25872 is transformable at similar levels to that of A1552, and as expected, transformation is PilT-dependent (Fig. 3B). However, in contrast to A1552, competence-induced cells lacking *pilT* did not aggregate and were excluded from aggregates formed by A1552 (Fig. 3C-E, H and I).

Given that all the other known components required for pilus assembly and transformation are highly conserved in ATCC25872, we tested whether PilA itself was responsible for this phenotype by exchanging the endogenous *pilA* open reading frame for that of A1552 (*pilA*ex). As expected, the ATCC25872 *pilA*ex strain was fully transformable (Fig. 3B). Importantly, however, in the absence of *pilT*, competence-induced cells of this strain were now able to form large aggregates, albeit at lower levels than in A1552 (Fig. 3C, F and G), and intermix within aggregates formed by A1552 (Fig. 3J and K).

### PilA variability governs auto-aggregation and enables kin-recognition

The results above suggest that the ability to aggregate is dependent on the particular PilA variant carried. Indeed, BLAST analyses of 647 *V. cholerae* genomes deposited in NCBI indicated that PilA exhibits extensive variation, whereas the other proteins encoded within its operon, as well as those from neighbouring genes, are all highly conserved (Fig. S9). Of the 636 intact *pilA* genes identified, the majority (492/636) encode a PilA identical to that of A1552, which is likely due to an over-representation of patient-derived pandemic strains in the database. Next, we extracted the unique PilA coding sequences (56/636) (Fig. S10A) and combined them with those of an in-house collection of various environmental and patient isolates. The resulting phylogenetic tree consists of around 12 distinct groups and was used to sample PilA diversity (Fig. 4A). As expected, the N-terminal ~50 residues, which contain sequences required for membrane trafficking and pre-pilin peptidase recognition, and that goes on to form the long structural alpha helix (a1), are well conserved (Fig. S10A and B). In contrast, apart from a few clusters of key conserved residues, including the C-terminal cysteines that form the characteristic disulphide-bonded loop, the C-terminal domains are highly variable (Fig. S10A and B).

**Figure 4.**
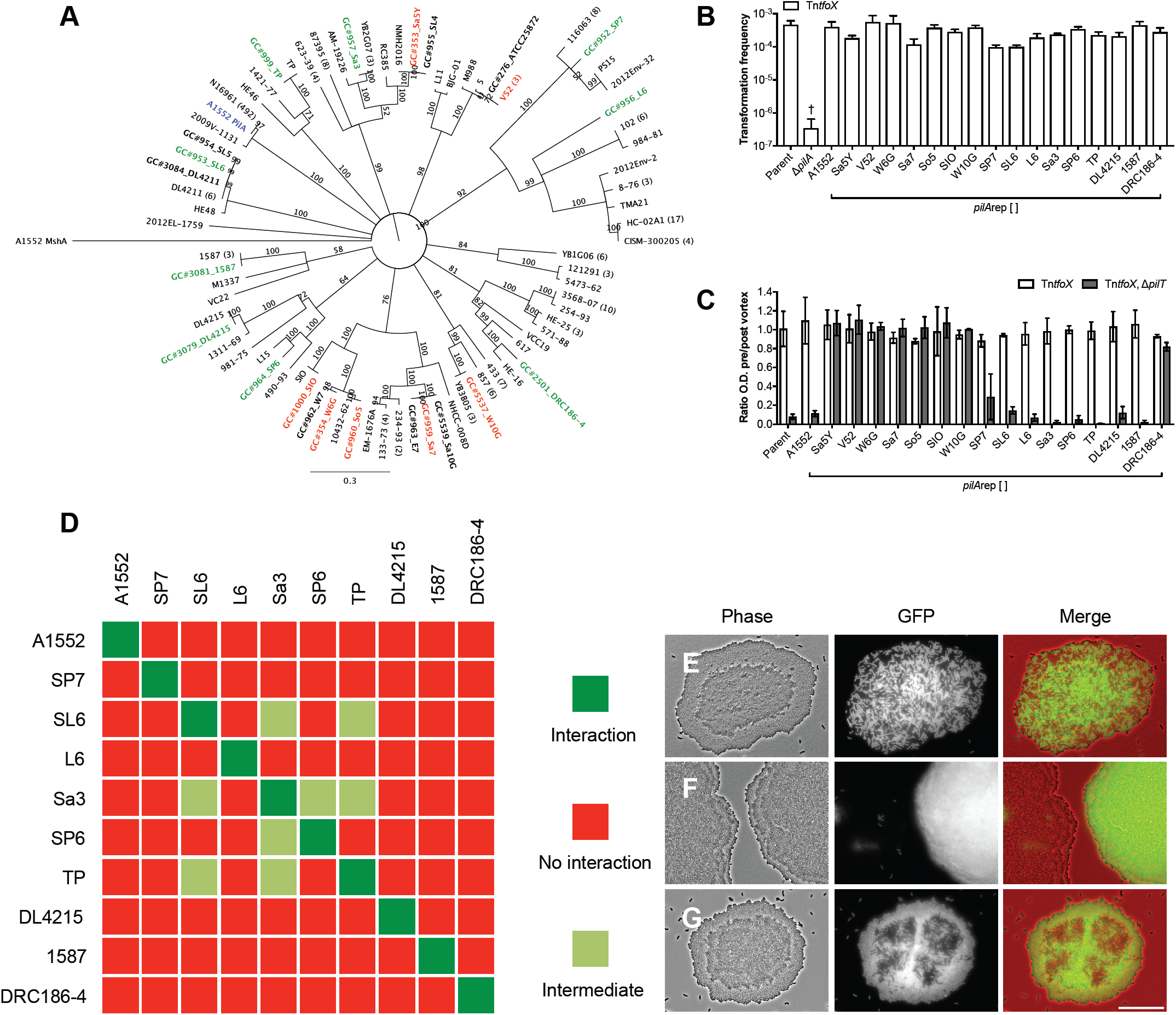
PilA variability governs auto-aggregation and enables kin-recognition. (**A**) Consensus neighbour-joining phylogenetic tree of PilA. The tree consists of the 56 unique PilA sequences identified from NCBI, 22 sequences from an in-house collection of various environmental and patient isolates (bold), and A1552 PilA (blue) and MshA (outgroup). Values shown are consensus support values (%). Aggregation capable (green) and incapable (red) PilAs tested in the *pilA*rep experiments are highlighted. Note that ATCC25872 and V52 PilA are identical. (**B-C**) Functionality of A1552-Tn*tfoX, pilA*rep[A1552] and 16 different PilAs assessed by transformation frequency and aggregation. (**B**) Chitin-independent transformation frequency assay. WT parent (A1552-Tn*tfoX*) and Δ*pilA* (A1552-Tn*tfoX*, Δ*pilA*) strains served as positive and negative controls. Transformation frequencies are the mean of three repeats (+S.D.). †, <d.l. in one repeat. (**C**) Aggregation was determined for the WT parent and each *pilA*rep strain in a *pilT*+ (A1552-Tn*tfoX*) and Δ*pilT* (A1552-Tn*tfoX*, Δ*pilT*) background, as indicated. Aggregation is shown as the ratio of the culture optical density (O.D. _600_) before and after vortexing, in the presence of *tfoX* induction. Values are the mean of three repeats (±S.D.). (**D**) Interaction matrix showing the results of a pairwise analysis of all possible interactions between aggregation proficient PilA. Interactions were tested by co-culturing non-fluorescent cells of the relevant Tn*tfoX*, Δ*pilT*, pilA*rep*[] strains with a fluorescent (GFP+) derivative of each strain. Cells were grown in the presence of *tfoX* induction. Self-self combinations served as controls. (**E-G**) Representative examples of (**E**) interaction, resulting in well-mixed aggregates (**F**) no interaction, resulting in fluorescent or non-fluorescent aggregates and (**G**) intermediate interactions, resulting in patterned aggregates. Cells were grown in the presence of *tfoX* induction. Merged images show GFP in green and phase-contrast (PC) in red. Bar = 25 μm.

To test how PilA variability affects pilus function but avoid the potential problems of working in various strain backgrounds, we inserted new *pilA* alleles at the native *pilA* locus in strain A1552, and used a short (30 bp) duplication of 3’ end of the original *pilA* to maintain any regulation of the downstream genes. We validated this *pilA* replacement (*pilA*rep) approach using A1552 PilA (*i.e.* Tn*tfoX, pilA*rep[A1552]), which is fully transformable, and in the absence of *pilT* aggregates at levels similar to the unmodified parent (Fig. 4B and C). We then tested 16 different PilA sequences from across the tree (Fig. 4A and Fig. S10B). Interestingly, all were equally capable of supporting transformation (Fig. 4B). In contrast, the ability to aggregate varied depending on the particular PilA. Indeed, in the absence of retraction, 9/16 PilAs supported the aggregation phenotype, though PilA DRC186-4 was intermediate, whereas 7/16 did not detectably aggregate (Fig. 4A and C).

Given that aggregation occurs via direct pilus-pilus interactions we hypothesised that the variability between the different aggregation-proficient PilA might allow pili composed of different PilA to distinguish between one another. To test this idea we used the same co-culture approach as before, using strains with and without constitutive GFP production, and examined all possible combinations (Fig. 4D). As expected, cells producing pili composed of an identical PilA always exhibited uniform mixing (Fig. 4D and E). Remarkably, however, in 41/45 possible unique combinations the interactions between different pili were highly specific (Fig. 4D). Indeed, cells of these strains aggregated in a pilin-specific manner, preferentially forming aggregates with cells producing pili composed of the same PilA. This resulted in aggregates that were either almost exclusively fluorescent or non-fluorescent (Fig. 4F). In 4/45 cases a partial cross-interaction was observed, resulting in an intermediate mixing phenotype, with aggregates composed of smaller but still segregated groups of cells (Fig. 4D and G). Overall these data indicate that pili composed of different PilA are able to discriminate between one another, probably via specific PilA-PilA interactions. Moreover, these data demonstrate that PilA variability not only determines the ability to aggregate, but also provides a mechanism for kin-recognition.

The TCP is also capable of aggregation under conditions mimicking virulence induction^18,57^. Indeed, in strains artificially induced for virulence TCP-mediated aggregates readily formed that appeared similar to those described in this work (Fig. S11A and C). As expected, they did not intermix with aggregates formed by DNA-uptake pili (Fig. S11B-D). Of note, however, is that TCP are largely limited to pandemic lineages with 2 variants of its major pilin TcpA: Classical and El Tor^58^-^60^. In contrast to the results above and in line with previous work^58^, a classical and El Tor strain formed uniformly mixed TCP aggregates (Fig. S11E), indicating that the different TcpA do not discriminate between one another.

### The unusual tail of ATCC25872/V52 PilA is an inbuilt inhibitor of aggregation

To rule out the possibility that strains unable to aggregate simply fail to make sufficient numbers of pili, we made labelable *pilA*rep cysteine variants of A1552 PilA (*pilA*rep[A1552; S67C]) and 2 non-aggregating alleles (*i.e. pilA*rep[Sa5Y; S67C] and *pilA*rep[V52; N67C]). All constructs behaved similarly to the equivalent non-cysteine variant, though the *pilA*rep[Sa5Y; S67C] has a modest transformation defect (Fig. S12A and B). Both the PilA[Sa5Y] and PilA[V52] cysteine variants were piliated at similar levels to the A1552 control and in the absence of *pilT* were hyper-piliated, although pili composed of PilA[Sa5Y] were generally very short (Fig. 5A and B). The hyper-piliation was especially clear for the V52 cysteine variant and provides direct evidence that the inability of some alleles to aggregate is likely not due to a failure to produce sufficient numbers of pili but reflects a specific property of the pilin itself. Notably, sheared pili of the *pilA*rep[A1552; S67C] variant assembled into large structures, as above, whereas those of the others did not (Fig. S12C).

**Figure 5.**
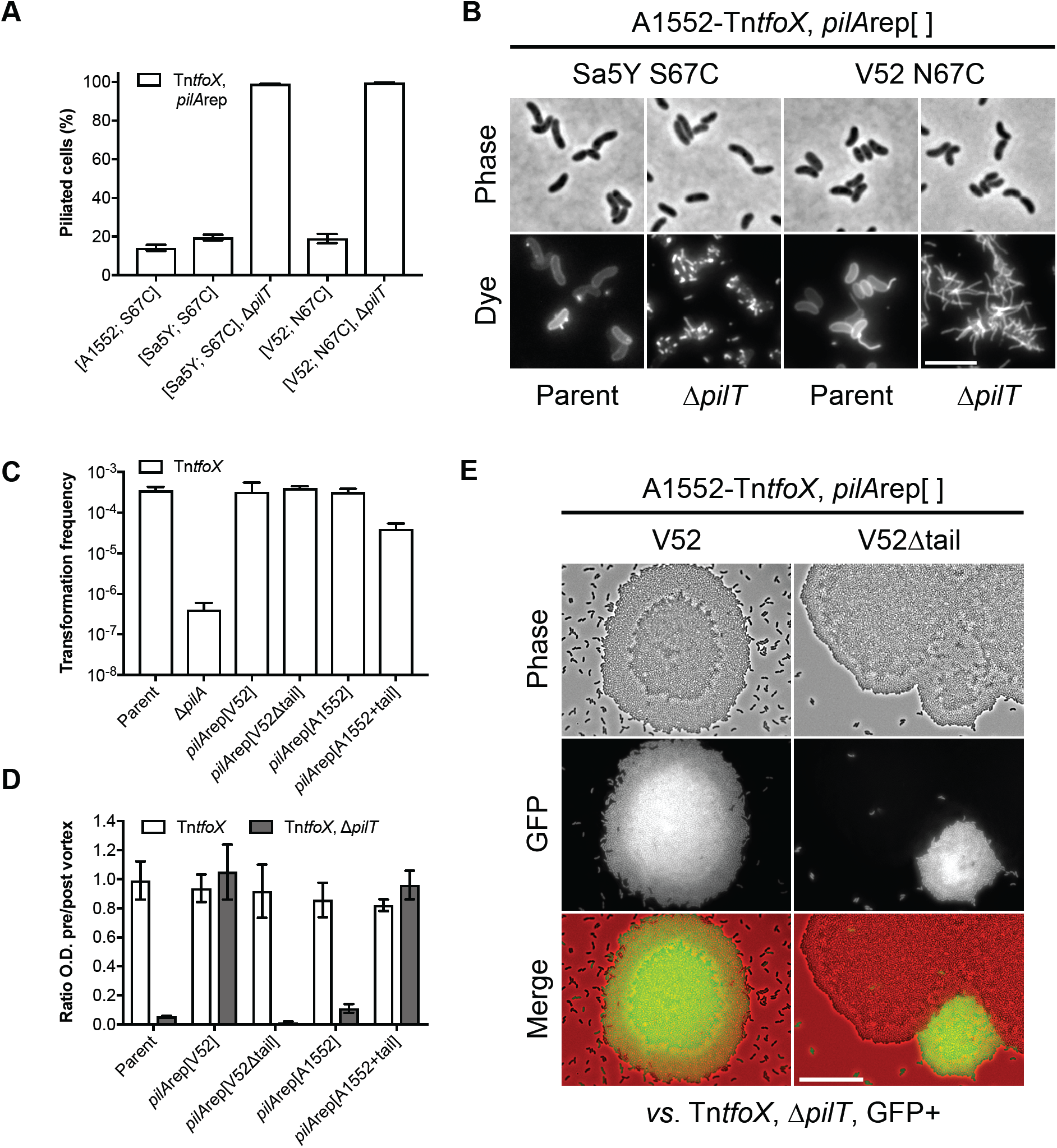
The unusual tail of ATCC25872/V52 PilA inhibits aggregation. (**A**) Quantification of piliation in snapshot imaging of cells of strains A1552-Tn*tfoX, pilA*rep[A1552; S67C], A1552-Tn*tfoX, pilA*rep[Sa5Y; S67C] and A1552-Tn*tfoX, pilA*rep[V52; N67C], in the indicated backgrounds. Cells were grown with *tfoX* induction and pili stained with AF-488-Mal. Bars represent the mean of three repeats (±S.D.). *n* = *c.a.* 2000 cells per strain per repeat. (**B**) Direct observation of pili stained with AF-488-Mal in cells carrying *pilA*rep[Sa5Y; S67C] and *pilA*rep[V52; N67C], in a WT parent (A1552-Tn*tfoX*) and Δ*pilT* (A1552-Tn*tfoX*, Δ*pilT*) background, as indicated. Bar = 5 μm. (**C-D**) Functionality of *pilA*rep tail variants assessed by natural transformation and aggregation. (**C**) Chitin-independent transformation assay. WT parent (A1552-Tn*tfoX*) and Δ*pilA* (A1552-Tn*tfoX*, Δ*pilA*) strains served as positive and negative controls. Transformation frequencies are the mean of three repeats (+S.D.). (**D**) Aggregation was determined for the WT parent and each *pilA*rep strain in a *pilT*+ (A1552-Tn*tfoX*) and Δ*pilT* (A1552-Tn*tfoX*, Δ*pilT*) background, as indicated. Aggregation is shown as the ratio of the culture optical density (O.D. _600_) before and after vortexing, in the presence of *tfoX* induction. Values are the mean of three repeats (±S.D.). (**E**) Co-culture of fluorescent cells of A1552-Tn*tfoX*, Δ*pilT*, GFP+, producing GFP, and non-fluorescent cells of either *pilA*rep[V52] (A1552-Tn*tfoX*, Δ*pilT, pilA*rep[V52]) or *pilA*rep[V52Δtail] (A1552-Tn*tfoX*, Δ*pilT, pilA*rep[V52Δtail]), as indicated. Note that ATCC25872 and V52 PilA are identical. Cells were grown in the presence of *tfoX* induction. Merged images show GFP in green and phase-contrast in red. Bar = 25 μm.

Strains V52 and ATCC25872 encode an identical PilA with an unusual repetitive C-terminal extension (SGSGSGSGSGSGSGSGSGN), or ‘tail’. Among the PilA sequences analysed here, there were 9 examples of this type of tail clustered into 2 well-separated phylogenetic groups (Fig. S8). Moreover, of the *pilA*rep PilAs unable to support aggregation 3/7 contain a tail (Fig. S8B). Thus, to test if the tail affected V52 PilA function we created a truncated variant *i.e. pilA*rep[V52Δtail], which was similarly transformable to the equivalent tailed version (Fig. 5C). Remarkably, however, removal of the tail restored the ability to aggregate in the absence of *pilT*, at levels indistinguishable from that of the PilA[A1552] controls (Fig. 5D), and demonstrated a similar ability to ‘recognise’ itself in co-culture experiments (Fig. 5E). These data suggest that the tail inhibits the ability of pili to aggregate, possibly by masking the site of pilus-pilus interaction. Indeed, a strain in which this tail was transplanted onto PilA of A1552 (*i.e. pilA*rep[A1552+tail] remained highly transformable but was unable to aggregate (Fig. 5C and D). However, since efforts to label this PilA variant have so far been unsuccessful we cannot exclude another, non-specific effect.

### DNA-uptake pili form networks on chitin surfaces

The data so far indicates that interactions between pili can mediate intercellular contacts. However, in liquid culture this relies on blocking pilus retraction. We reasoned that if these interactions are relevant to the normal ecology of *V. cholerae* then we might be able to detect them under more realistic conditions. Therefore, we visualised pili produced by otherwise WT cells upon cultivation on chitinous surfaces (Fig. 6A). Strikingly, under these conditions of natural induction, cells colonising the chitin surface were generally found embedded within dense networks of pili that were often overlaid by larger pili structures (Fig. 6A). In contrast to the chitin-independent liquid culture experiments, cells on the chitin surface often appeared to possess multiple pili simultaneously (Fig. 6B), though the crowded nature of the cells precluded direct quantification. Furthermore, since the washing steps required for pilus staining tend to remove cells, but not pili, from the surface, the number of cells engaged in these networks is likely underestimated. Control experiments confirmed that labelling on surfaces is specific and that these structures are DNA-uptake pili (Fig. S13A-C). Notably, Δ*pilT* cells are heavily hyper-piliated under these natural induction conditions (Fig. S13D), indicating that the phenotypes observed in liquid culture are not an artefact of artificial competence induction.

**Figure 6.**
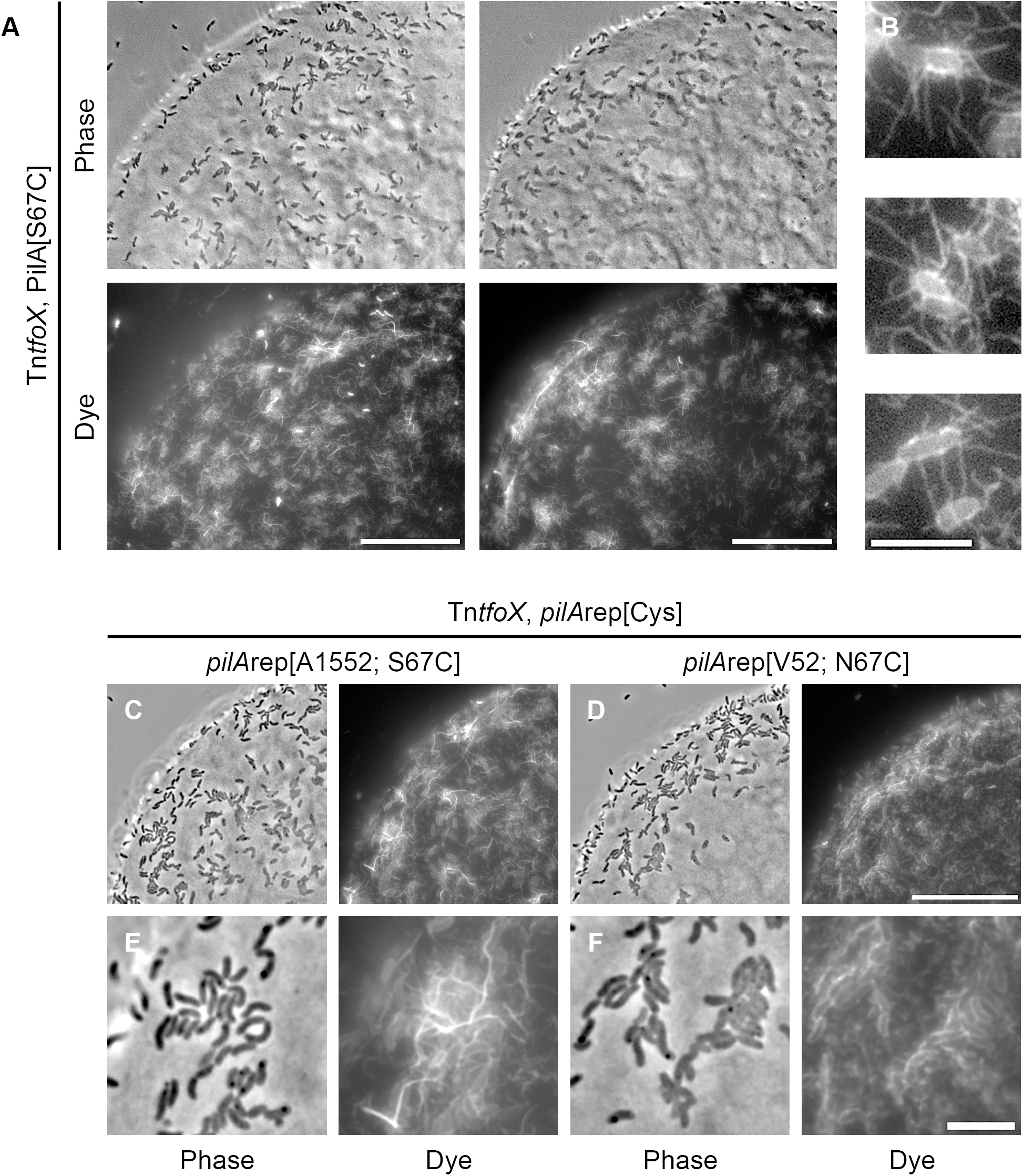
DNA-uptake pili form networks on chitin surfaces. (**A-B**) Chitin beads were stained with AF-488-Mal after incubation for 48h in defined artificial seawater (DASW) with cells of A1552-Tn*tfoX*, PilA[S67C], as indicated. The figure depicts two separate examples of colonised surfaces, bars = 25 μm. (**B**) Insets show enlarged examples of cells producing multiple pili. Bar = 5 μm. (**C-F**) Chitin beads were stained with AF-488-Mal after incubation for 48h in DASW with cells of either (**C**) A1552-Tn*tfoX, pilA*rep[A1552; S67C] or (**D**) A1552-Tn*tfoX, pilA*rep[V52; N67C], as indicated. Bar = 25 μm. Panels (**E** and **F**) show enlargements of the surfaces shown in (**C** and **D**). Note the absence of large pili networks in (**F**). Bar = 5 μm.

Next, using the *pilA*rep cysteine variants described above, we directly compared the pili assembled on chitin surfaces by cells producing the aggregation-proficient PilA[A1552] with those producing the aggregation-deficient PilA[V52]. Consistent with its functionality in the various assays above, cells carrying *pilA*rep[A1552; S67C] were also found within similar pili networks (Fig. 6C and E). In contrast, such networks were never observed for cells carrying *pilA*rep[V52; N67C] (Fig. 6D and F). Instead, the surfaces were coated in what appeared to be mostly individual pili. This difference was particularly clear at later time-points when cells presumably began to exhaust the chitin surface. Cells producing PilA[A1552] covered surfaces with extensive and long-range networks of pili that were maintained within detached pieces of biofilm (Fig. S14A and C). In contrast, for cells producing PilA[V52] the surfaces were coated with individual pili and never large networks (Fig. S14B and D). In summary, these data suggest that the aggregation phenotypes observed in liquid culture experiments report the natural ability of pili to interact on chitin surfaces.

## Discussion

Here we demonstrate that DNA-uptake pili are highly dynamic, that these dynamics are PilT-dependent, and that cells lacking *pilT* are minimally transformable, providing direct evidence for the longstanding model, whereby pilus retraction facilitates DNA-uptake. Indeed, our results on pilus dynamics are in close agreement with those recently reported by Ellison *et al.*, who notably, went on to demonstrate that the pilus binds directly to DNA^61^. The major finding of this work, however, is that DNA-uptake pili are able to interact and distinguish between one another in a specific manner depending on their sequence. In liquid culture when retraction is blocked this manifests as an exaggerated auto-aggregation phenotype. Since only a subpopulation of cells is piliated at any one time, and these pili are dynamic, blocking retraction likely facilitates auto-aggregation by producing a homogenous population of hyper-piliated cells, thereby increasing the chances of interactions between pili. Work in *Neisseria meningitidis*, which auto-aggregates naturally at low levels but is dramatically enhanced by the deletion of *pilT*, supports this idea^62,63^. Importantly, however, under natural induction conditions on chitin surfaces, cells producing pili capable of aggregation elaborate multiple pili and form dense pili networks in an otherwise unmodified background, indicating that the chitin surface likely promotes interactions between pili. The crowded surface environment might inherently foster these interactions. Alternatively, the altered physiology of cells growing on chitin might also impact pilus assembly via effects on the extension/retraction motors.

The TCP of *V. cholerae* is induced during virulence and is essential for colonisation^18,57,64^. Under laboratory conditions TCP production results in auto-aggregation similar to that described here^17,19,58^. Moreover, in an infection model cells on the intestinal cell surface were found encased within networks of TCP, that were suggested to protect cells from host defences^65^. Given the similarities between the two systems, especially our observation of dense pili networks on colonised chitin surfaces, we propose that the DNA-uptake pilus might play an analogous role in the aquatic environment. Indeed, as hypothesised elsewhere^29,66^, since pilus production is dependent on an intact chitin-utilisation pathway, colonisation mechanisms using DNA-uptake pili would be inherently selective for (i) nutritious chitinous surfaces and (ii) favour the recruitment and retention of productive cells while excluding non-productive cells unable to make pili. Thus, intercellular interactions between DNA-uptake pili could act at multiple stages of colonisation, but might be particularly advantageous during early stages to bring smaller chitin particles together, and thus provide resistance to protozoan grazers, as well as at later stages to keep cells together during biofilm dispersal. It remains possible, however, that these interactions represent an ancient colonisation mechanism that has since been replaced. Arguing against this idea is the observation that all pandemic strains retain the same interaction-proficient PilA even though other environmental strains have been free to vary their PilA without impacting its ability to mediate transformation. Interestingly, ingestion of colonised chitin particles is thought to facilitate transmission to humans^26^ and recovering cholera patients exhibit a strong immune response to PilA^67^. One possibility that should be investigated is that the networks of DNA-uptake pili we observed on chitin surfaces protect cells during this process.

In *Neisseria gonorrhoeae*, artificially varying the density or post-translation modification state of pili leads to a form of cell sorting based on differential interaction forces between pili^68^. This effect is likely related to aggregate dispersal during its infective lifestyle but does not permit specific recognition *per se*^69^. In contrast, the discovery here that the natural variability of PilA controls the ability of pili to self-interact, and creates highly specific interactions, provides a direct mechanism for kin recognition. The best-studied examples of kin recognition in microorganisms all involve adhesins and some form of aggregation (*e.g.* Flo1; *Saccharomyces cerevisiae*^70^, TgrB1-TgrC1; *Dictyostelium discoideum*^71^, and TraA; Myxobacteria^72^). In evolutionary terms, these recognition mechanisms are classified as ‘greenbeards’ because the cue, recognition of the cue and the resulting cooperative activity are all encoded by the same gene^73,74^. The ability of DNA-uptake pili to recognise and interact with pili composed of the same kind of PilA fits this classification and is therefore a specific form of greenbeard recognition^73,74^. However, this form of recognition implies close identity only at the greenbeard locus and so is better referred to as kind recognition^73,74^.

An important question going forward will be to understand what drives PilA diversity and how this is related to the type VI secretion system, which acts to kill non-kin bacteria^75,76^. Similarly, the apparent acquisition of an inhibitor of pilus interactions by some PilA (*e.g.* ATCC25872/V52) hints that the ability to interact may not always be beneficial. Therefore, future work should focus on how the pili networks we observed on chitin surfaces contribute to the ecology of *V. cholerae*, especially under environmental conditions. Indeed, we still know relatively little about the natural lifestyle of *V. cholerae* on chitin, in part due to the inherent technical difficulties associated with manipulating these surfaces. Nevertheless, the demonstration that in liquid media, specific interactions between pili composed of different major pilins is sufficient to enable segregation, provides a robust proof-of-concept that T4P have the ability to function as a recognition mechanism. Finally, the fact that (i) T4P are widespread, (ii) auto-aggregation via T4P has been reported in multiple species^77^ and (iii) the major pilin subunit often varies^4,52^, raises the possibility that specific interactions between T4P might be quite common and therefore represent an important contribution to bacterial kin recognition worthy of continued investigation.

## Materials and Methods

### Bacterial strains and plasmids

The bacterial strains used in this study are shown in Supplementary Table S1, together with the plasmids used and their construction. A1552, the *V. cholerae* strain used throughout this work^78^, is a toxigenic O1 El Tor Inaba strain representative of the on-going 7^th^ cholera pandemic, and was derived from a traveller entering the United States after being infected on a commercial aeroplane that took off in Peru^79^.

### General methods

Bacterial cultures were grown aerobically at 30°C or 37°C, as required. Liquid medium used for growing bacterial strains was Lysogeny Broth (LB-Miller; 10 g/L NaCl, Carl Roth, Switzerland) and solid medium was LB agar. Where indicated, LB-S contained 20 g/L NaCl. Ampicillin (Amp; 100 μg/mL), gentamicin (Gent; 50 μg/mL), kanamycin (Kan; 75 μg/mL), streptomycin (Str; 100 μg/mL) and rifampicin (Rif; 100 μg/mL) were used for selection in *E. coli* and *V. cholerae*, as required. To induce expression from the *P*_BAD_ promoter, cultures were grown in media supplemented with 0.2% L-arabinose. Natural transformation of *V. cholerae* on chitin flakes was done in 0.5x DASW (Defined artificial seawater), supplemented with vitamins (MEM, Gibco) and 50 mM HEPES, as previously described^30^. Counter-selection of phenylalanyl-tRNA synthetase (*pheS*\*) insertions (Trans2 method; see below) was done on medium supplemented with 20 mM 4-chloro-phenylalanine (cPhe; Sigma-Aldrich, Switzerland). Thiosulfate citrate bile salts sucrose (TCBS; Sigma-Aldrich, Switzerland) agar was used to counter-select for *E. coli* following bacterial mating. SacB-based counter-selection was done on NaCl-free medium containing 10 % sucrose.

### Strain construction

DNA manipulations and *E. coli* transformations were carried out using standard methods^80^, and all constructs were verified by PCR and Sanger sequencing (Microsynth AG, Switzerland). Genetic engineering of *V. cholerae* was done using a combination of natural transformation and FLP-recombination; Trans-FLP^81,82^, *pheS*\*-based counter-selection; Trans2^83,84^, and allelic exchange using bi-parental mating and the counter-selectable plasmid pGP704-Sac28^29^. The mini-Tn7 transposon carrying *araC* and various *P*_BAD_-driven genes was integrated into the large chromosome by tri-parental mating, as previously described^85^.

### Chitin-independent competence induction

Chitin oligosaccharides resulting from growth on chitin trigger natural competence induction via the production of a master regulator, TfoX^29,30,86-88^. High cell-density, as sensed by quorum sensing via HapR, results in the production of an intermediate regulator QstR, which acts in concert with TfoX to regulate the transcription of a subset of the competence genes^30,41,89^. Therefore, to induce natural competence in liquid culture we used a well characterised and already validated chitin-independent approach that results in low levels of TfoX production^41^. This approach is based on the integration of a mini-Tn7 transposon into the large chromosome of *V. cholerae* containing an arabinose-inducible copy of *tfoX* (*i.e. araC, P*_BAD_-*tfoX*), which we refer to as Tn*tfoX*. In the presence of inducer, strains carrying Tn*tfoX* turn on the expression of the competence genes according to the known regulatory pathways and upon reaching high cell-density are transformable at levels similar to those seen on chitin^41^. In the absence of inducer, competence genes are not produced and strains are non-transformable^41^.

### Transformation frequency assay

Diverse strains harbouring Tn*tfoX* were tested for transformation using a chitin-independent transformation frequency assay, as previously described^32,41^. Briefly, overnight cultures were back-diluted 1:100 in fresh media with and without arabinose, as indicated, and grown 3h at 30°C with shaking (180 rpm). 0.5 mL aliquots of the cultures were mixed with 1 μg genomic DNA (GC#135; A1552-lacZ-Kan) in 1.5 mL eppendorf tubes and incubated 5h at 30°C with shaking (180 rpm), prior to serial dilution in PBS (Phosphate buffered saline) and enumeration after overnight growth on LB media in the absence and presence of kanamycin. Transformation frequency was calculated as the number of transformants divided by the total number of bacteria.

### Pilus shearing assay

Cultures were grown for 6h at 30°C with shaking (180 rpm) in 25 mL LB + 0.2% arabinose within a 125 mL Erlenmeyer flask. To shear pili from the cell surface, 10 mL culture was removed, vortexed at max speed for 1 min, and cells removed by three sequential centrifugation steps (10min; 4000 x *g*; 4°C). To the resulting supernatant saturated ammonium sulphate was added to 40% and incubated on ice for 1h. Precipitated proteins were recovered by centrifugation (30min; 20,000 x *g*; 4°C) and washed once with PBS. Samples were then re-suspended in 2x Laemmli buffer, boiled (15min; 95°C) and stored at  −20°C until needed. To compare PilA levels between samples the re-suspension volume was normalised according to the optical density of the starting culture. Total protein controls were intact cell lysates. The relative amount of PilA in each sample was determined by Western blotting.

### Aggregation assay

Overnight cultures were back-diluted 1:100 in the absence and presence of arabinose, as needed, and grown in 14 mL round bottom polystyrene test tubes (Falcon, Corning) on a carousel style rotary wheel (40 rpm) at 30°C. After 6h growth, aggregates were allowed to settle by standing the tube at RT for 30 min. The optical density at 600 nm (O.D._600_) of the culture was then measured before and after mechanical disruption (vortex max speed; ~5 sec), which served to disperse any settled aggregates and return them to solution. Aggregation is expressed as the ratio of the O.D._600_ Pre/Post-vortexing. For time-course experiments the standing time was reduced to 5 min. To visualise aggregates by microscopy overnight cultures were back-diluted either 1:100 individually or 1:200 when mixed, as needed, and were grown for 4h, as described above.

### Microscopy

Cells were mounted on microscope slides coated with a thin agarose pad (1.2% w/v in PBS), covered with a #1 cover-slip, and were observed using a Zeiss Axio Imager M2 epifluorescence microscope attached to an AxioCam MRm camera and controlled by Zeiss AxioVision software. Image acquisition was done using a Plan-Apochromat 100×/1.4 Ph3 oil objective illuminated by an HXP120 lamp. Images were analysed and prepared for publication using ImageJ (http://rsb.info.nih.gov/ij).

### Pilus staining and quantification

Alexa Fluor™ 488 C_5_ Maleimide (AF-488-Mal; Thermo Fisher Scientific; Cat# A10254) was dissolved in DMSO, aliquoted and stored at −20°C protected from light. Cultures used for staining were grown for 3.5h at 30°C on the rotary wheel, as above, in the absence and presence of competence induction, as required. To stain cells 100 μL of culture was mixed with dye at a final concentration of 25 μg/mL^39^ and incubated at RT for 5 min in the dark. Stained cells were harvested by centrifugation (5000 x *g*; 1 min), washed once with LB, re-suspended in 200 μL LB and imaged immediately. For quantification of piliation in snapshot imaging approximately 2000 cells per strain were analysed in each of three independent repeats. A subset of this data set was also analysed for the number of pili per cell and pilus length, as indicated in the text. For time-lapse analysis, images were acquired at RT at 10 sec intervals for 1 min (*i.e.* 7 frames). To analyse the number of pili produced per cell, per min, 5 fields of cells were analysed for each of the three independent repeats, yielding an analysis of 1947 cells in total.

### Imaging on chitin surfaces

Visualisation of pilus production on chitin surfaces was done using chitin beads (NEB; Cat#S6651), which have previously been validated as a useful analogue for the natural chitin surface^41,75^. Prior to use, chitin beads were washed 5x with 0.5x DASW + 50 mM HEPES + vitamins. Colonisation of chitin beads was done in 12-well culture plates (Cellstar) by mixing 0.1mL washed O/N culture with 0.1 mL washed beads in a final volume of 1 mL 0.5x DASW + 50 mM HEPES + vitamins, and incubated at 30°C; 48h. Pilus staining was done as described above except that the final re-suspension volume was reduced to 50 μL. Stained beads were mounted directly on glass microscope slides, covered with a #1 cover-slip and imaged using either a Plan-Apochromat 100x/1.4 Ph3 or Plan-Neofluar 40x/1.3 Ph3 oil objective, as needed. To image cells after prolonged incubation (≥72h), plates were incubated within a homemade humidified chamber. To avoid damaging the beads manipulations were done using wide bore tips.

### Western blotting

Cell lysates were prepared by suspending harvested cells in 2x Laemmli buffer (100 μL buffer per O.D. unit) before boiling at 95°C for 15 min. Proteins were separated by SDS-PAGE using a 15% resolving gel and blotted onto PVDF membranes using a wet-transfer apparatus. Immuno-detection was performed as described previously^41^. Primary anti-PilA antibodies were raised in rabbits against synthetic peptides of A1552 PilA (Eurogentec, Belgium; #1510525) and used at a dilution of 1:5000. Anti-Rabbit IgG HRP (Sigma; Cat# A9169) diluted 1:5000 was used as a secondary antibody. Sample loading was verified with anti-RNA Sigma70-HRP (BioLegend; Cat# 663205) diluted 1:10,000.

### Motility assay

To quantity motility phenotypes 2 μL overnight culture was spotted soft LB agar (0.3%) plates (two technical replicates) and incubated at RT for 24h prior to photography. The swarming diameter (cm) was measured and is expressed as the mean of three independent biological repeats. A flagellin-deficient (Δ*flaA*) non-motile strain was used as a negative control.

### Bioinformatics of PilA diversity

*Vibrio cholerae* genomes were obtained from NCBI (National Center for Biotechnology Information), and are listed in Supplementary File S1. Geneious software (10.2.3)^90^ was used to perform custom BLAST analyses and identify *pilA*. Unique PilA sequences were extracted and combined with the PilA sequences from strain A1552 and a collection of environmental isolates, as deduced by Sanger sequencing (Supplementary File S2), as indicated in the text. PilA sequences were aligned with Muscle and a consensus neighbour-joining tree constructed using the Jukes-Cantor substitution model, resampled with 100 bootstrap replicates. MshA from strain A1552 was used as an outgroup.

## Reproducibility

All data shown are representative of the results of at least three independent biological repeats.

## Acknowledgements

We thank Ivan Mateus-Gonzalez for assistance with bioinformatics analyses. We further thank A. Boehm, S. Pukatzki, J. Mekalanos, and members of the Institut National de Recherche Biomédicale of the Democratic Republic of the Congo, for providing *V. cholerae* strains and V. Pelicic for advice on *pheS*-mediated counter-selection. Work on this problem was supported by a Marie Sklodowska-Curie Individual Fellowship (703340; CMDNAUP) to D.W.A. and by EPFL intramural funding and a Starting Grant from the European Research Council (ERC; 309064-VIR4ENV) to MB. M.B. is a Howard Hughes Medical Institute (HHMI) International Research Scholar (Grant# 55008726).

## Author contributions

Conception, design and analysis: D.W.A and M.B. Performed research: D.W.A, S.S, C.S and M.B Wrote the manuscript: D.W.A and M.B.

## Declaration of Interests

The authors declare no competing interests.

